# Intact Somatosensory Temporal Sensitivity in Adults on the Autism Spectrum: A High-Density Electrophysiological Mapping Study Using the Mismatch Negativity (MMN) Sensory Memory Paradigm

**DOI:** 10.1101/2024.02.05.578908

**Authors:** Emily L. Isenstein, Edward G. Freedman, Sophie Molholm, John J. Foxe

## Abstract

Atypical reactivity to somatosensory inputs is common in autism spectrum disorder and carries considerable impact on downstream social communication and quality of life. While behavioral and survey work have established differences in the perception of somatosensory information, little has been done to elucidate the underlying neurophysiological processes that drive these characteristics. Here, we implemented a duration-based somatosensory mismatch negativity paradigm to examine the role of temporal sensitivity and sensory memory in the processing of vibrotactile information in autistic (n=30) and neurotypical (n=30) adults. To capture the variability in responses between groups across a range of duration discrepancies, we compared the electrophysiological responses to frequent standard vibrations (100 ms) and four infrequent deviant vibrations (115, 130, 145, and 160 ms). The same stimuli were used in a follow-up behavioral task to determine active detection of the infrequent vibrations. We found no differences between the two groups with regard to discrimination between standard and deviant vibrations, demonstrating comparable neurologic and behavioral temporal somatosensory perception. However, exploratory analyses yielded subtle differences in amplitude at the N1 and P220 time points. Together, these results indicate that the temporal mechanisms of somatosensory discrimination are conserved in adults on the autism spectrum, though more general somatosensory processing may be affected. We discuss these findings in the broader context of the MMN literature in autism, as well as the potential role of cortical maturity in somatosensory mechanisms.

## INTRODUCTION

Atypical sensory reactivity, including in response to tactile input, is commonly reported in autism spectrum disorder (ASD, American Psychiatric Association, 2013). As the first sensory system to fully develop (Piontelli, 2015), touch is vital during early childhood development (Blackwell, 2000; Cascio et al., 2019; Rheingold, 1966) and provides feedback on the relationship between the self and the physical world (Riemann & Lephart, 2002). While research on somatosensation in autism has increased in recent years (Buyuktaskin et al., 2021; Noda et al., 2022; Shafer et al., 2021), the great majority of studies have focused on auditory and visual perception, despite evidence suggesting that over sixty percent of people with autism have higher tactile sensitivity than their neurotypical (NT) peers (Tomchek & Dunn, 2007). These alterations in sensory processing can affect quality of life and may have downstream effects on higher-order features such as social communication and repetitive behaviors that are characteristic in autism (Kawakami & Otsuka, 2021). For example, avoidance of caregiver touch at 9 months has been associated with parental reports of autistic behaviors at 18 months (Mammen et al., 2015), and both over-responsiveness (Cascio et al., 2016) and under-responsiveness (Foss-Feig et al., 2012) to touch in children with autism have been associated with differences in social behavior and communication.

Atypical tactile processing also presents inherent challenges to everyday life. Pain and discomfort can result from hyper*-*sensitivity to touch, which is characterized by disproportionate, negative responses to tactile stimuli perceived as neutral by most individuals (Baranek et al., 1997). Tactile hypo-sensitivity can also be found in autistic people, causing slow or absent reactions to harmful sensations such as burns or blisters, often compounded by the fact that these individuals do not seek out care (Jussila et al., 2020). This variability in somatosensory processing is found across the autism spectrum, but little is known about what drives these sensitivities. Largely, findings have been inconsistent, with many showing equivalent tactile detection thresholds between neurotypical and autistic groups (Cascio et al., 2008; Güçlü et al., 2007; O’Riordan & Passetti, 2006). Others, however, have found differences depending on the qualities of the stimuli, such as amplitude (Puts et al., 2013), frequency (Blakemore et al., 2006), or whether the stimulation was static or dynamic (Puts et al., 2014). Given the various types of receptors and pathways used to detect different types of somatosensation – including pressure, vibration, temperature, and more – it remains challenging to establish whether there is a common alteration along the sensory processing pathway that drives atypical somatosensory processing in autism.

Despite the accumulation of behavioral evidence for dysregulation of the sense of touch in autism and its apparent role in foundational features of autism, relatively few studies have examined the neurophysiology of tactile perception in autistic individuals. Some have found comparable electrophysiological responses between autistic and neurotypical groups (Butler et al., 2017; Dwyer et al., 2022; Kadlaskar et al., 2021), while others have shown differentiation in latency (Espenhahn et al., 2021; Hashimoto et al., 1986), hemispheric lateralization (Miyazaki et al., 2007), and response amplitude (Russo et al., 2010). Differences also emerge when the task involves active engagement (Deschrijver et al., 2017; Kemner et al., 1994). Atypical modulations of somatosensory event related potentials (SEPs) by emotion (Fanghella et al., 2022) and visual attention (Noda et al., 2022) have also been recorded. Dissociable electrophysiological correlates associated with tactile hyper- and hypo- responsiveness suggest that this variability in tactile sensitivity may be indexed at distinct points in the sensory-perceptual pathway (Azouz et al., 2014; Cascio et al., 2015).

Temporal processing has been identified as another possible driver of autism features, potentially contributing to cognitive differences such as difficulties with joint attention or behavioral effects such as sensory overload (Allman, 2011). Differential temporal cognition has been identified in the tactile domain in autism (Ide et al., 2019; Wada et al., 2014); for example, people with autism who demonstrated greater temporal resolution had higher reported hypersensitivity to sensory stimuli, and greater sensitivity to tactile stimuli correlated with restricted and repetitive behaviors and interests (Ide et al., 2019). An extended window of time in which stimuli from two different senses are bound together and presumed to come from the same source - called the multisensory temporal binding window - has also been suggested in the visual-tactile domain (Greenfield et al., 2015; Ropar et al., 2018), yet little has been done to probe temporal sensitivity within the somatosensory system in autism using direct neurophysiological assays of brain function.

One means of examining temporal processing is through the use of the well-established mismatch negativity (MMN) paradigm. In the typical MMN experimental design, identical stimuli (standards) are presented repeatedly in a sequence with a small fraction of the stimuli (deviants) differing from the majority along a specific feature dimension (e.g. frequency, location, or duration) (Molholm et al., 2005; Näätänen et al., 1989; Sams et al., 1985). Under such conditions, there is a distinct electrophysiological signal associated with deviant detection. A considerable bolus of prior work has established that MMN generation to deviant features - such as the duration change used herein - occurs pre-attentively and does not require active attentional engagement with the stimulus stream (Alho et al., 1992; Näätänen et al., 1980; Novak et al., 1992; Ritter et al., 2006). Accordingly, study participants can engage with other inputs - such as watching a movie or reading - during an MMN experiment, allowing for its deployment as a neural measure in neurodevelopmental conditions where attentional functioning may be atypical or disordered.

The MMN can be elicited across multiple sensory domains (Butler et al., 2012; Butler et al., 2011; Joutsiniemi et al., 1998; Kekoni et al., 1997; Näätänen et al., 2007; Ritter et al., 2002; Tales et al., 1999), but limited work has been conducted using the somatosensory MMN (sMMN), particularly in autism. In an oddball paradigm that utilized an auditory standard, an auditory oddball, and a novel tactile stimulus, no difference in response to the tactile input was found between children with autism and controls (Kadlaskar et al., 2021). The only other related study in autism that we are aware of found no evidence for a MMN effect when the location of stimulus presentation distinguished standards from deviants (Kemner et al., 1994). As such, there is a gap in knowledge on whether autism affects recognition of deviant somatosensory inputs. Here, we were specifically interested in the recognition of temporal discrepancies in the somatosensory domain, since as outlined above, prior work has pointed to a potential association between heightened tactile temporal sensitivity and clinical features of autism (Ide et al., 2019).

To this end, we employed a sMMN paradigm that measured the electrophysiological response to standard vibrations (100 ms) and four deviant vibrations of elongated duration (115 ms, 130 ms, 145 ms and 160 ms). By varying the extent of duration deviance across different experimental blocks, we aimed to test the sensitivity of the tactile sensory memory system to duration differences that ranged from quite small to relatively large. Our prior work using an identical paradigm in neurotypical adults showed that this span of duration deviants adequately captures the range of possible MMN responses, from essentially undetectable (115 ms deviant) to fully detectable (160 ms), and gradations in between (Isenstein et al., 2023). Following the electrophysiological part of the experiment, during which participants were not informed about the presence or purpose of the deviant vibrations (i.e. participants watched silent videos), a subsequent behavioral test was conducted to determine if participants could actively detect the various deviant durations when they were explicitly attended to. Finally, we assessed whether electrophysiological and behavioral sMMN metrics were associated with cognitive and behavioral measures to assess the clinical relevance of potential atypical somatosensory temporal processing.

## METHODS

### Participants

Sixty-two adults between the ages of 18 and 45 were recruited, 31 with a diagnosis of autism and 31 non-autistic individuals. One individual in the autism group was excluded due to poor data quality and one in the non-autism group was excluded due to clinical suspicion of autistic traits. All participants had self-reported normal hearing, normal or corrected to normal vision, and none reported a history of traumatic brain injury or psychosis-related disorder. Individuals in the neurotypical group (n=30, mean age: 25.60, STD: 5.97; 18M/12F (1 non-binary)) were recruited from the local Rochester area and had no first-degree relatives with an autism diagnosis. Data from fifteen of the thirty individuals in the neurotypical group were included in previously published work using the identical experimental design (Isenstein et al., 2023). People in the autism group (n=30, mean age: 25.95, STD: 7.31; 18M/12F (2 non-binary)) were recruited from the local Rochester area and research databases, and all had a previous diagnosis of autism made by a clinician that was confirmed using the Autism Diagnostic Observation Schedule 2 (ADOS-2) (Lord et al., 2012) by a research-reliable administrator. Participants in the autism group taking stimulant medication were asked to withhold these medicines on the day of the EEG visit; no participants in the neurotypical group took stimulant medication. No participants in either group took medication for psychosis-related conditions. All participants were administered the Wechsler Abbreviated Scale of Intelligence (Wechsler, 2011), and participants were not excluded based on IQ results. The following assessments were also completed by each participant or by a legal guardian, as appropriate: the Social Responsiveness Scale – (SRS-2) (Constantino & Gruber, 2012), the Repetitive Behavior Scale – Revised (Bodfish et al., 2000), the Adolescent/Adult Sensory Profile (Brown & Dunn, 2002) and the Adult ADHD Self-Report Screening Scale for DSM-5 (ASRS-5) (Ustun et al., 2017).

Of note, the adults in the autism group were informally surveyed on whether they prefer person first (person with autism) or identity first language (autistic person). The cohort was split relatively evenly between these two options, as well as ‘no preference’; accordingly, both person and identity first language are used in this paper.

All participants, or caregivers as appropriate for adults with assigned legal guardians, provided written informed consent. Adults with assigned legal guardians also provided verbal and written assent to participate. All procedures were reviewed and approved by the Research Subjects Review Board (RSRB) at the University of Rochester (STUDY00002036), and conformed with the tenets for ethical conduct of human subjects’ research laid out in the Helsinki declaration (“World Medical Association Declaration of Helsinki: ethical principles for medical research involving human subjects,” 2013). All participants were paid an hourly rate for their participation.

### EEG Stimuli and Task

The EEG task parameters used here were identical to those described in our previous work that characterized parametric duration somatosensory mismatch negativity in neurotypical adults (Isenstein et al., 2023), and followed procedures developed for auditory duration MMN mapping (De Sanctis et al., 2009). An Adafruit 1201 vibrating mini-motor disc (Arduino, Turin, Italy) was secured around the participant’s right index fingertip with a strip of hook and loop fasteners to administer somatosensory stimulation, with confirmation that the disc was snug but not uncomfortably tight against the finger pad. The mini-motor disc administered vibrations at a frequency of 183.33 Hz, produced at 5 volts via Arduino Uno microcontroller that was operated through Presentation® software (Version 18.0, Neurobehavioral Systems, Inc., Berkeley, CA). Participants sat in an electrically-shielded and sound-attenuating EEG booth (IAC Acoustics, North Aurora, IL, USA) and wore ear protection (MPOW HM035A; Longgang, Guangdong, China) to preclude air-conducted auditory input produced by the vibrating mini-motor disc. Participants watched a silent video of their choosing and were instructed to ignore the vibrotactile stimuli while attending to the video. They were also told to keep their right hand still with their palm facing up so the mini-motor disc would not make contact with the table and alter the vibration. Vibrations were presented at a ratio of 80% standard duration (100 ms) and 20% deviant duration (115, 130, 145, or 160 ms) across four randomly ordered condition blocks. Each block contained a total of 1000 stimuli (inter-stimulus interval of 750 ms±150 ms) in a pseudo-random order such that no deviant vibrations occurred consecutively. Participants were not informed that there would be any variation in the vibration types until the behavioral portion.

High-density EEG was recorded using a 128-channel BioSemi EEG system (Biosemi ActiveTwo, Amsterdam, The Netherlands) with a DC to 150 Hz pass-band and referenced to the common mode sense (CMS) active electrode. Data were recorded continuously at 512 Hz and the default Biosemi decimation filter of 1/5 of this rate was applied prior to processing using the EEGLAB toolbox (version 14.1.2) (Delorme & Makeig, 2004) through MATLAB 2021a software (Mathworks, Natick, MA, USA). As with the prior study using this paradigm (Isenstein et al., 2023), data were of high quality in the unfiltered state and the data were not further filtered prior to the remaining pre-processing steps. Channel interpolation was conducted on channels with amplitudes ± 3 standard deviations from the mean. The average reference was used and automatic artifact rejection was implemented based on the normal distribution of maximum and minimum amplitudes. Data were epoched from -100 to 500 ms with the zero-point set at the onset of the vibration, then baseline corrected to the 100 ms prior to vibration onset. Trials were rejected if they contained any amplitudes of +/- 200 µV. The average of a cluster of five frontal electrodes surrounding the Fz scalp site (see **Figure 6D**) comprised the region of interest for the MMN measures derived herein (as outlined in our prior work (Isenstein et al., 2023) and previous literature (Chen et al., 2014; Kekoni et al., 1997; Spackman et al., 2007)). The “topoplot” function in the EEGLAB toolbox was used to produce topographic maps that represent voltage patterns mapped onto a 2-dimensional circular representation of the scalp.

### Behavioral Discrimination Task

Participants also completed a brief behavioral discrimination task to assess perceptual thresholds following the EEG session. This behavioral task had the same stimulus ratio but included only 100 total stimuli per block. Stimuli were divided in four blocks, one for each of the deviant durations, and counter-balanced across participants, each containing 80 standard and 20 deviant stimuli. Unlike the electrophysiological paradigm, participants were directed to attend to the somatosensory stimuli and indicate when they detected an infrequent longer stimulus by pressing a button with their left hand. No video was presented during this task. One participant in the autism group was unable to complete the behavioral task.

### Data Analyses

Data were analyzed in the same manner as in Isenstein et al., 2023. The latency of the MMN for each individual participant was calculated by finding the point of maximum difference between their standard and deviant waveforms between 150 and 250 ms post-stimulus. The individual latencies were averaged on a group level yielding the following mean values for the neurotypical group: 115-dur = 183 ms; 130-dur = 182 ms; 145-dur = 183 ms; 160-dur = 192 ms, for an average of 185 ms. The autism group yielded the following values: 115-dur = 200 ms; 130-dur = 186 ms; 145-dur = 190 ms; 160-dur = 204 ms, for an average of 195 ms. The average amplitude during the 50 ms window surrounding the neurotypical mean of 185 ms (160-210 ms) was used as the metric of the MMN amplitude in both groups.

A mixed-design analysis of variance (ANOVA) was used to compare the latency in the MMN time window using a within-subjects factor of DEVIANCE LEVEL (115, 130, 145, 160 ms) and a between-subjects factor of GROUP. Mixed design ANOVA was also used to compare amplitude in the MMN time window using within-subjects factors of DEVIANCE LEVEL (115, 130, 145, 160 ms) and DURATION (Standard vs. Deviant) and a between-subjects factor of GROUP. Difference waves were calculated by subtracting the individual standard waveforms from each respective deviant waveform for visualization purposes.

D-prime values (d’) were also calculated as normalized hit rates minus normalized false alarm rates (Green & Swets, 1966). This value served as a metric of sensitivity to behaviorally detect the deviant vibrations and compared between groups using a mixed ANOVA with a within-subjects factor of DEVIANCE LEVEL (115, 130, 145, 160 ms) and a between-subjects factor of GROUP.

Pearson correlations were conducted between experimental measures (each of the four D-prime values) and clinical measures (SP Touch Processing (2 missing from each group), SRS-Total (1 missing from autism group), RBS-R Self Injury (1 missing from each group), and ASRS -5 Total (1 missing from neurotypical group, 2 missing from autism group)). *P*-values were corrected for multiple comparisons using the Benjamini-Hochberg method with a False Discovery Rate of 0.05 (Benjamini & Hochberg, 1995); because of the conservative nature of this correction, we still report the uncorrected results.

### Post-Hoc Exploratory Analysis (Statistical Cluster Plots – SCP)

In a follow-up exploratory analysis phase, we further probed this rich high-density ERP dataset using the statistical cluster plot (SCP) approach (Molholm et al., 2002). This *post-hoc* method is intended as a follow-up to the pre-planned *a-priori* tests of the sMMN described above, purely as a means to better describe the recorded data. Given the hypothesis-free nature of this approach, any additional effects uncovered must be treated as *post-hoc* and should be interpreted cautiously. Rather, such effects should be considered hypothesis generation tools and would need to be confirmed/replicated in a follow-up study.

SCP maps were generated comparing Standard and Deviant amplitudes within groups (see **Figure 5**) using a cluster-based permutation test whereby a series of two-tailed, paired sample t-tests (critical and cluster α = 0.05) were calculated across all timepoints and all electrode sites using the FieldTrip toolbox in Matlab. Monte-Carlo sampling with 5000 iterations allowed for modeling of significance probability while the triangulation method was used for spatial clustering with correction for multiple comparisons.

To further compare components of the ERPs in response to Standard vibrations between groups, all of the Standard trials were averaged together for each group (henceforth called ‘All Standard’). Cluster-based permutation testing using the same methodology as above (with the exception that independent sample t-tests were used instead of paired sample t-tests) were also calculated to temporally and spatially compare the ‘All Standard’ responses between groups.

Two time periods of interest were identified for further investigation. In the ‘All Standard’ waveform at the same frontal cluster of electrodes, the minimum negative amplitude between 50 and 150 ms in the neurotypical group was centered at 108 ms and the average amplitude during the 50 ms window surrounding 108 ms (83-133 ms) was calculated for both groups and is henceforth called the N1. Additionally, the maximal positive amplitude between 150 and 250 ms in the neurotypical group was centered at 224 ms and the average amplitude of the 50 ms window surrounding 225 ms (199-249 ms) was calculated for both groups and is henceforth called the P220. Independent samples t-test were used to compare the ‘All Standard’ N1 and P220 amplitudes between groups.

Pearson correlations were conducted between experimental measures (‘All Standard’ N1 and P220 amplitudes) and clinical measures (Touch Processing (2 missing from each group), SRS-Total (1 missing from autism group), RBS-R Self Injury (1 missing from each group), and ASRS -5 Total (1 missing from neurotypical group, 2 missing from autism group)). Benjamini-Hochberg False Discovery Rate was again used for correction for multiple comparisons.

## RESULTS

### Demographic data

Chi Square analysis did not identify a significant difference in sex among groups [X^2^ (1, n = 60) = 0.00, p = 1.00]. The groups did not differ by age (t(58) = -0.32, *p* = 0.75) or IQ (t(56) = -0.49, *p* = 0.63). The groups differed significantly on all selected clinical measures, including SP Touch Processing (t(54) = -3.52, *p* = 8.98*10^-4^), SRS-2 Total (t(57) = -7.69, *p* = 2.30*10^-10^), RBS-R Self Injury (t(55) = -2.53, *p* = 0.01), and ASRS -5 Total (t(55) = -3.92, *p* = 2.51*10^-4^). See **Table 1** for group average demographic data.

**Table 1.** Demographic data for each group.

|  | Amplitude Group Mean (SE) |  | p value |
| --- | --- | --- | --- |
|  | Neurotypical | Autism |  |
| Sex | 18 M/ 12 F | 18 M/ 12 F | 1.00 |
| Age | 25.49 (1.10) | 25.95 (1.33) | 0.75 |
| Race |  |  |  |
| White | 14 | 25 |  |
| Black or African American | 2 | 1 |  |
| Asian | 12 | 3 |  |
| Hawaiian or Pacific Islander | 0 | 0 |  |
| American Indian | 0 | 0 |  |
| Multiple Races | 2 | 1 |  |
| Unknown/Not reported | 0 | 0 |  |
| Ethnicity |  |  |  |
| Hispanic or Latino | 2 | 2 |  |
| Not Hispanic or Latino | 26 | 26 |  |
| Unknown/Not Reported | 2 | 2 |  |
| WASI (Full Scale) | 110.79 (2.95) | 108.62 (3.29) | 0.63 |
| SP – Touch Processing | 27.64 (1.19) | 34.07 (1.39) | $8.98^{-4}$ |
| SRS – Total Score | 41.93 (3.87) | 92.90 (5.42) | $2.30^{-10}$ |
| RBS-R – Self Injurious Behavior | 1.79 (0.84) | 4.62 (0.74) | 0.01 |
| ASRS-5 Total Score | 8.86 (0.66) | 13.03 (0.83) | $2.51^{-4}$ |
| *A-Priori Results* |  |  |  |

### Behavioral Data

The behavioral detection sensitivity (D-prime) values for each of the Deviance Levels for both groups are shown in **Table 2** and plots of the distribution of D-prime in both groups can be found in **Figure 1**.

**Figure 1.**
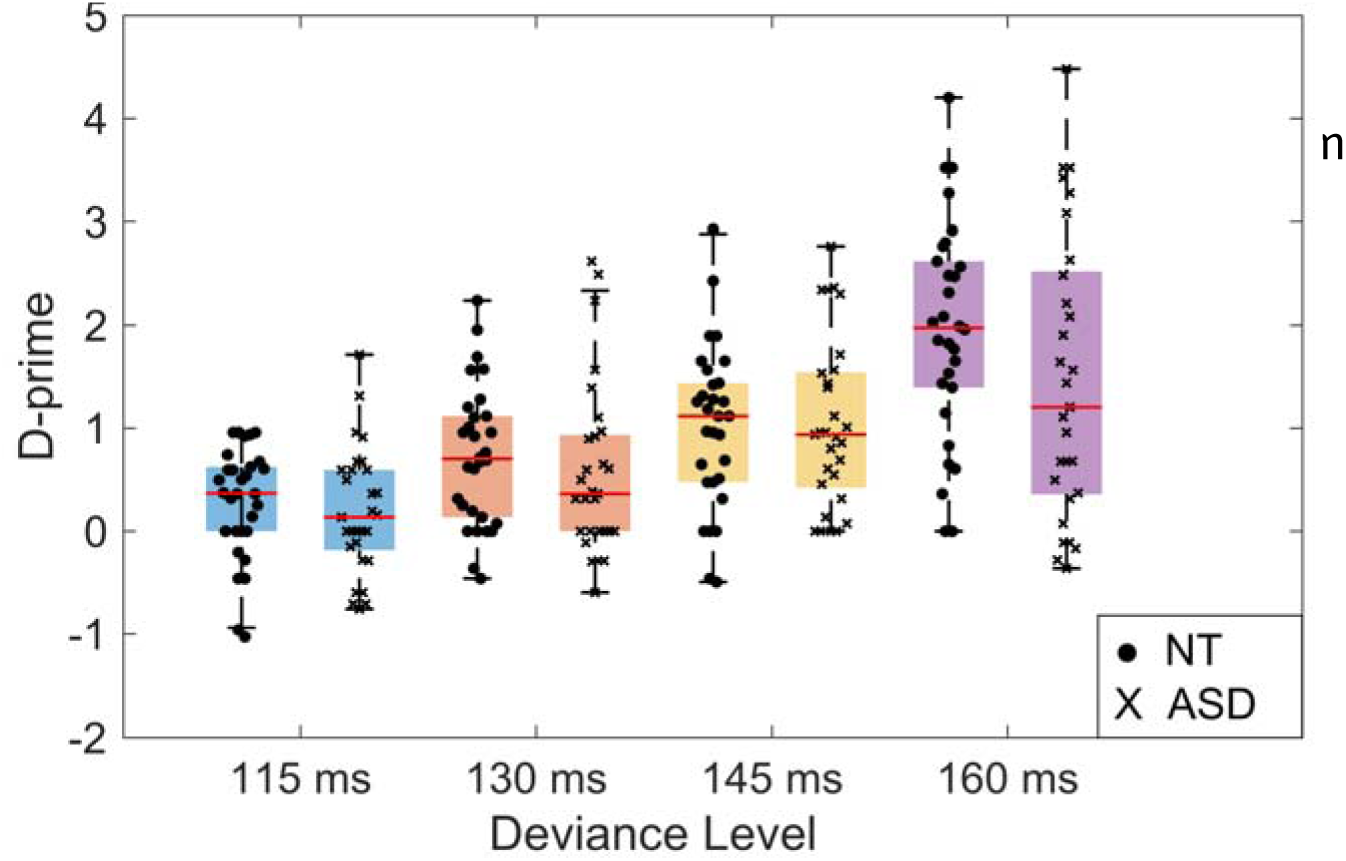
Behavioral sensitivity measures. Individual participant D-prime values are shown for each duration condition in the neurotypical (●) and autism (x) groups.

**Table 2.**
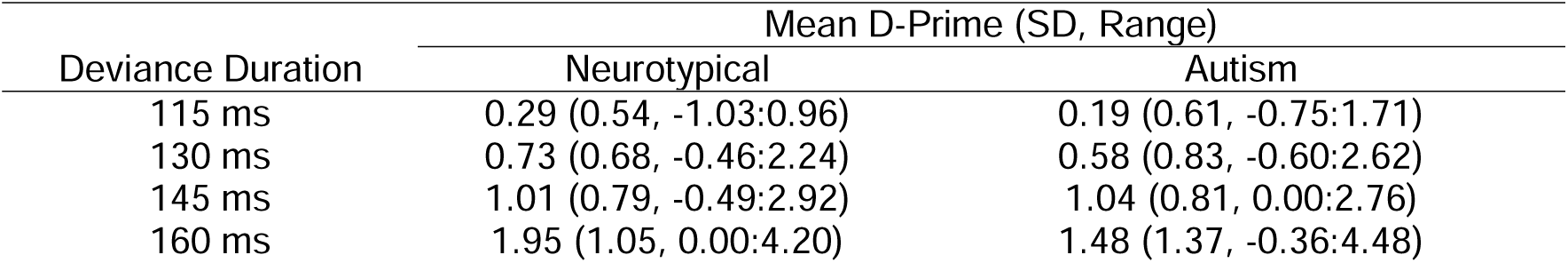
Mean D-Prime values for the Neurotypical and Autism groups at each Deviance Duration.

Detection of the deviant vibrations was quite difficult, and some participants in each group did not correctly detect any deviant stimuli at certain Deviance Levels (115 dev: 4 NT, 5 ASD; 130 dev: 4 NT, 6 ASD; 145 dev: 3 NT, 4 ASD; 160 dev: 2 NT, 0 ASD). The mixed-design ANOVA yielded a significant main effect of DEVIANCE LEVEL (F(3, 171) = 50.81, *p* = 1.57*10^-23^, η^2^ = 0.47), reflecting an increase in D-prime in both groups as the extent of deviance increased. There was no interaction between DEVIANCE LEVEL and GROUP (F(3, 171) = 1.49, *p* = 0.22, η^2^ = 0.03) and no main effect of group (F(3, 171) = 1.08, *p* = 0.30, η = 0.02). Sphericity was not violated (*p* = 0.07).

Given the main effect of DEVIANCE LEVEL, follow-up protected pairwise comparisons were conducted to unpack this effect; means and standard errors are reported. These revealed that the 115 ms d’ (x□ = 0.24±0.08) was significantly smaller than the 130 ms d’ (x□ = 0.65±0.10, *p* = 8.10*10^-4^, 95% C.I. = -0.65, -0.18), 145 ms d’ (x□ = 1.03±0.11, *p* = 1.25*10^-9^, 95% C.I. = -1.00, -0.57) and the 160 ms d’ (x□ = 1.72±0.16, *p* = 1.36*10^-14^, 95% C.I. = -1.77, -1.19). The 130 ms d’ was significantly smaller than the 145 ms d’ (*p* = 0.01, 95% C.I. = -.60, -0.15) and the 160 ms d’ (*p* = 5.73*10^-10^, 95% C.I. = -1.35, -0.78), and the 145 ms d’ was significantly smaller than the 160 ms d’ (*p* = 1.85*10^-7^, 95% C.I. = -.92, -0.45).

### MMN Latency

With regard to latency measures of the MMN, mixed-design ANOVA yielded no significant interaction between DEVIANCE LEVEL and GROUP (F(3,174) = 0.88 , *p* = 0.44, η = 0.02), but there was a main effect of DEVIANCE LEVEL (F(3,174) = 4.99 , *p* = 4.40*10^-3^, η = 0.08) and a main effect of GROUP (F(1,58) = 7.04, *p* = 0.01, η = 0.11). Sphericity was violated (*p* = 0.01). Per the main effect of DEVIANCE LEVEL, protected pairwise comparisons showed that the peak latency to the 160 ms deviant condition in the MMN window (x□ = 201.27±: 3.16) was significantly longer than the that of the 130 ms (x□ = 183.85±2.83, *p* = 1.32*10^-4^, C.I. = 8.96, 26.08) and 145 ms (x□ = 186.65±2.81, *p* = 3.63*10^-4^, C.I. = 6.94, 22.50) deviant latencies. All other *p*’s > 0.08. The neurotypical group (x□ = 186.47±2.30) also had a significantly shorter latency than the autism group (x□ = 195.12±2.30, *p* = 0.01, C.I. = -15.17, -2.12).

### MMN Amplitude

With regard to ERP amplitudes within the MMN time window, mixed-design ANOVA yielded an interaction between DURATION and DEVIANCE LEVEL (F(3, 174) = 17.98, *p* = 3.30*10^-10^, η^2^ = 0.24). There was no interaction between DURATION and GROUP (F(1, 58) = 0.71, *p* = 0.40, η^2^ = 0.01), DEVIANCE LEVEL and GROUP (F(3, 174) = 1.23, *p* = 0.30, η = 0.02) and DURATION, DEVIANCE LEVEL, and GROUP (F(3, 174) = 0.65, *p* = 0.59, η^2^ = 0.01). There was a significant main effect of DURATION (F(1, 58) = 138.87, *p* = 5.01*10^-17^, η^2^ = 0.71) and DEVIANCE LEVEL (F(3, 174) = 6.63, *p* = 2.98*10^-4,^ η^2^ = 0.10), as well as a between-subjects effect of GROUP (F(1, 58) = 4.72; *p* = 0.03, η^2^ = 0.08). Sphericity was not violated (DEVIANCE LEVEL *p* = 0.14, DURATION * DEVIANCE LEVEL *p* = 0.32). Waveforms are plotted in **Figure 2** and difference waves are plotted in **Figure 3** and group means can be found in **Table 3**.

**Figure 2.**
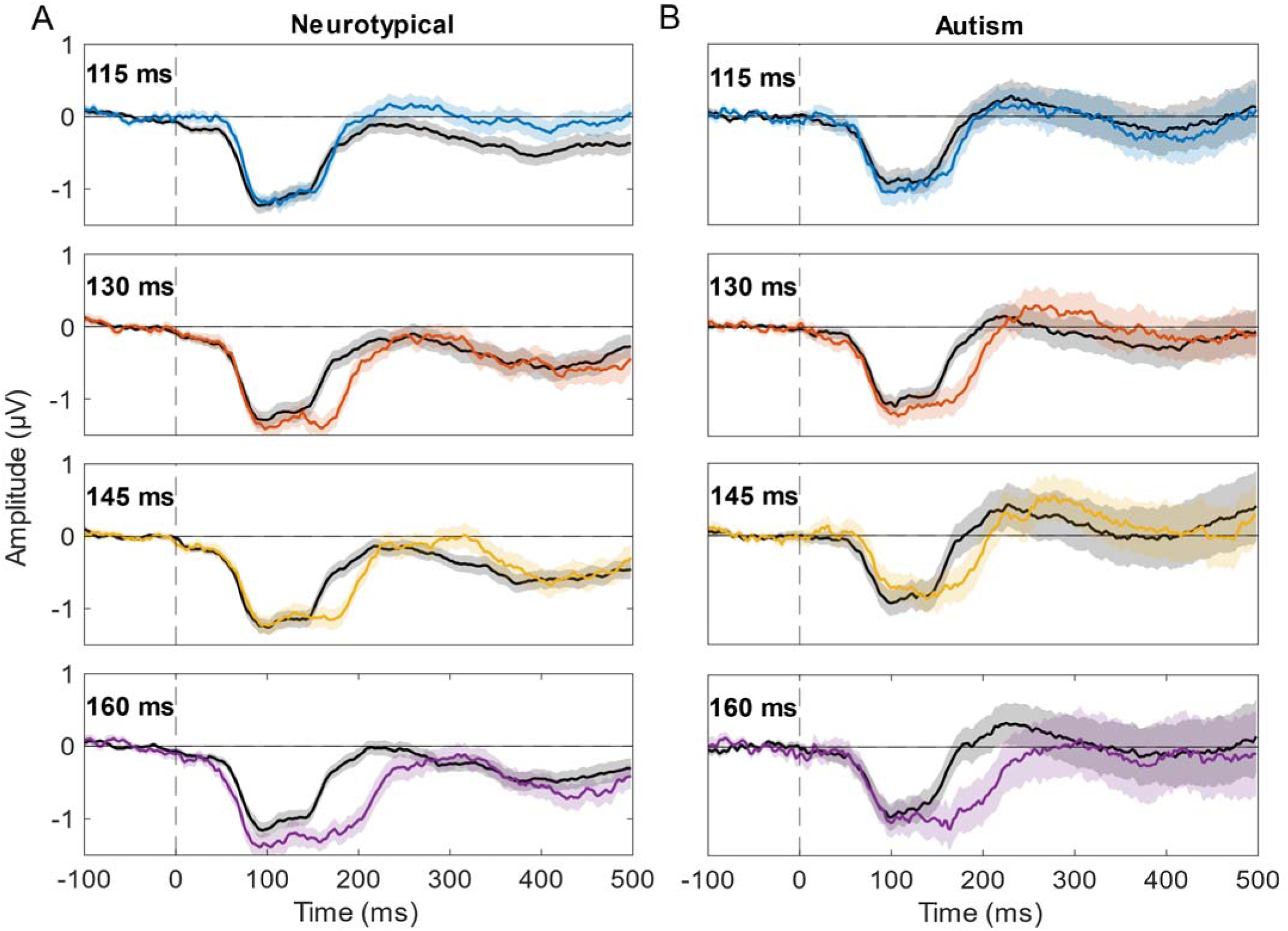
Group-averaged standard versus deviant responses for each of the four MMN conditions. **A)** MMN responses to each deviant condition in the neurotypical group. **B)** MMN responses to each deviant condition in the autism group. Waveforms to the standard 100 ms tone are plotted in black. Standard error of the mean is indicated by shading around the waveforms.

**Figure 3.**
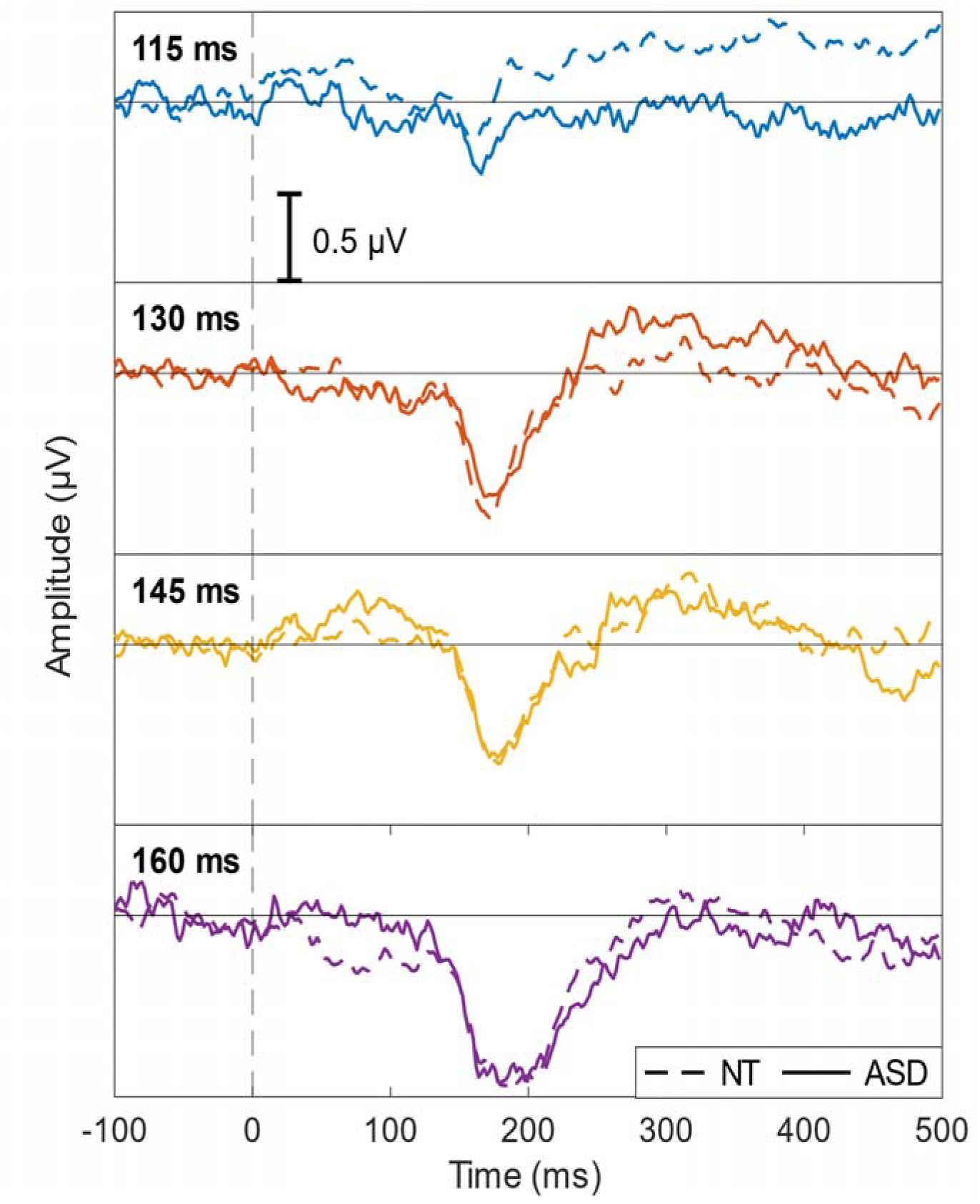
Group comparisons of the derived MMN waveforms. Difference wave plots comparing the neurotypical (dotted lines) and autism (solid lines) differences in response between the standard and each of the deviant conditions.

**Table 3.** Mean amplitudes of 115, 130, 145 and 160 ms deviant MMNs (Deviant – Standard) in Neurotypical and Autism Groups.

|  | Amplitude Group Mean (SE) |  |
| --- | --- | --- |
|  | Neurotypical | Autism |
| MMN |  |  |
| 115 Dev - Std | 0.08 µV (0.11) | -0.17 µV (0.11) |
| 130 Dev - Std | -0.51 µV (0.08) | -0.52 µV (0.13) |
| 145 Dev - Std | -0.47 µV (0.11) | -0.48 µV (0.13) |
| 160 Dev - Std | -0.86 µV (0.10) | -0.86 µV (0.11) |

Follow-up protected paired comparisons were employed to unpack the significant main effects and interactions revealed by the mixed-design ANOVA. The Deviant amplitudes (x□ = -0.67±0.07) were significantly more negative than the Standard amplitudes (x□ = -0.20±0.07, *p* = 5.01*10^-17^, C.I. = -0.55, -0.39). With regard to DEVIANCE LEVEL, amplitudes in the 115 ms condition (x□ = -0.24±0.08) were significantly smaller than the amplitudes in the 130 (x□ = -0.54±0.08, *p* = 2.03*10^-4^, 95% C.I. = 0.15, 0.45), 145 (x□ = -0.43±0.10, *p* = 0.03, 95% C.I. = 0.03, 0.35), 160 ms (x□ = -0.55±0.08, *p* =4.40*10^-5^, 95% C.I. = 0.17, 0.45) conditions. The autism group (x□ = -***waveforms.*** *Difference wave plots comparing the neurotypical (dotted lines) and autism (solid lines) differences in response between the standard and each of the deviant conditions.* 0.29±0.10) had overall lower magnitude amplitudes in the MMN time window than the neurotypical group (x□ = -0.58±0.10, *p* = 0.03, 95% C.I. = -0.56, -0.02). All other *p*’s > 0.10. It is important to point out that this effect is with respect to all vibrations regardless of DURATION or DEVIANCE LEVEL, and thus does not reflect processes specifically associated with the MMN.

### Topography

Topographic plots were created to visualize the MMN difference waves (Deviant minus Standard) across the scalp at the critical time-points selected for the MMN (100, 140, 180, and 220 ms) (**Figure 4**). The results recapitulate the above statistics, with comparable plots between the neurotypical and autism groups at all time points and conditions. Dipolar fields are evident in the 130, 145, and 160 ms deviant conditions over left/central scalp, echoing the plots in Figure 2.

**Figure 4.**
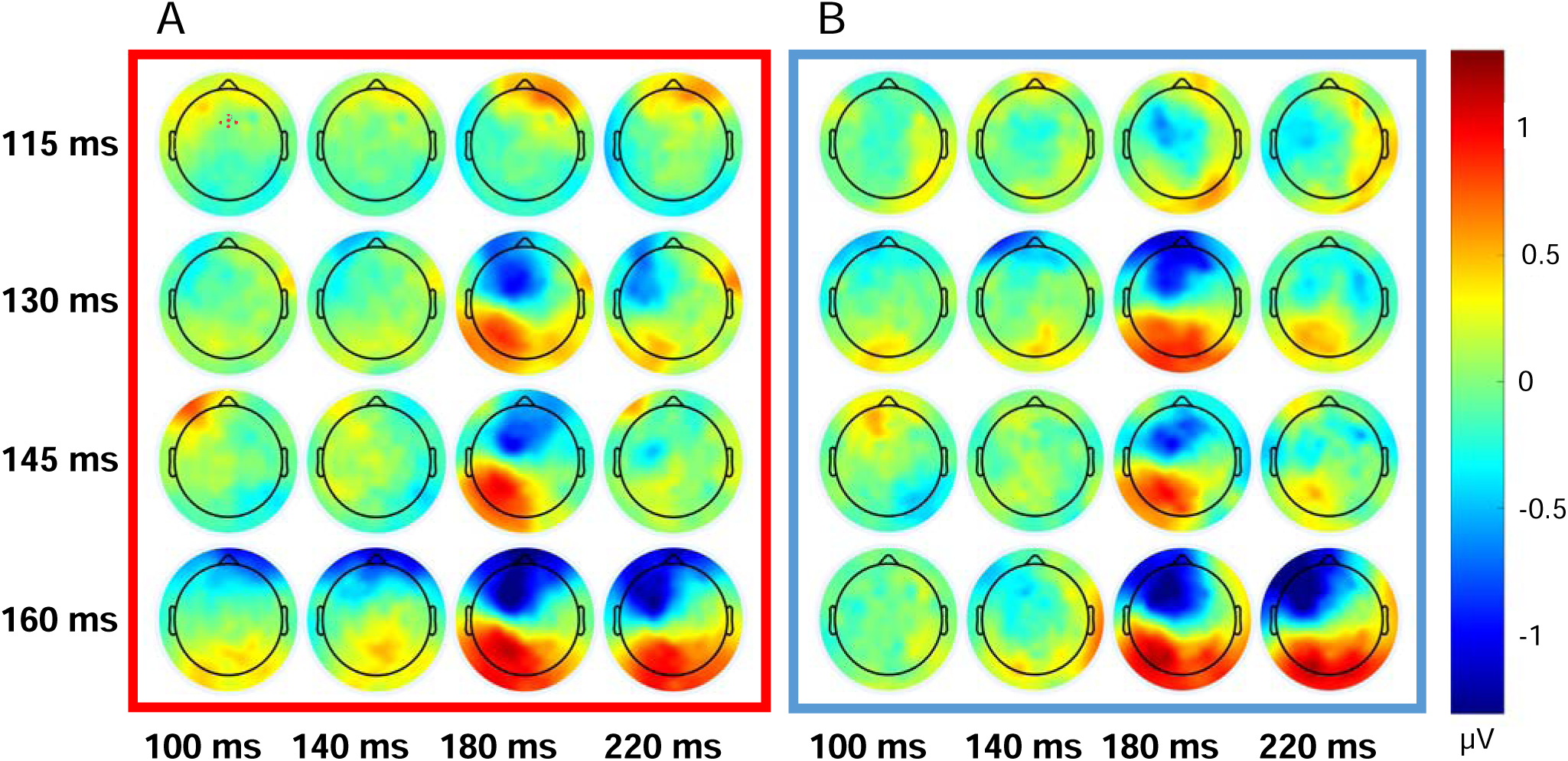
Visualization of dipolar activity across several timepoints surrounding the MMN. Topographic potential plots of the MMN difference waves in the period between 100 and 220 ms poststimulus for each of the 115 ms, 130 ms, 145 ms, and 160 ms deviant conditions in **A)** neurotypical and **B)** autism groups. The specific frontal electrodes used in the statistical analyses are highlighted in upper left plot of **A**.

### Correlations

Pearson correlations between experimental and clinical measures yielded significant uncorrected correlations between the RBS-R Self-Injurious Behavior Score and both the 130 (r = -0.33, *p_uncorrected_* = 0.01, *p_corrected_* = 0.16) and 160 ms (r = -0.30, *p_uncorrected_* = 0.02, *p_corrected_* = 0.18) D-prime. Neither survived correction for multiple comparisons. All other uncorrected *p*’s > 0.29. See **Supplemental Figure 1A,B** for pertinent correlation plots. Since these tests did not survive correction, they are not discussed further here.

#### Post-hoc Results

### Cluster Plots

**Figure 5** shows statistical cluster plots that visualize the scalp positions and time points during which the standard and deviant conditions significantly differed from each other. These plots emphasize the strong similarities between the neurotypical and autism groups across conditions. As found in Isenstein et al., the 130, 145, and 160 ms deviant conditions demonstrate synchronized regions of significant MMN that was not present in the 115 ms deviant, now shown in both groups (Isenstein et al., 2023). Specifically, the 150-250 ms post-stimulus time window captures the MMN, with the significant clusters most pronounced over the fronto-central and central-parietal scalp regions.

**Figure 5.**
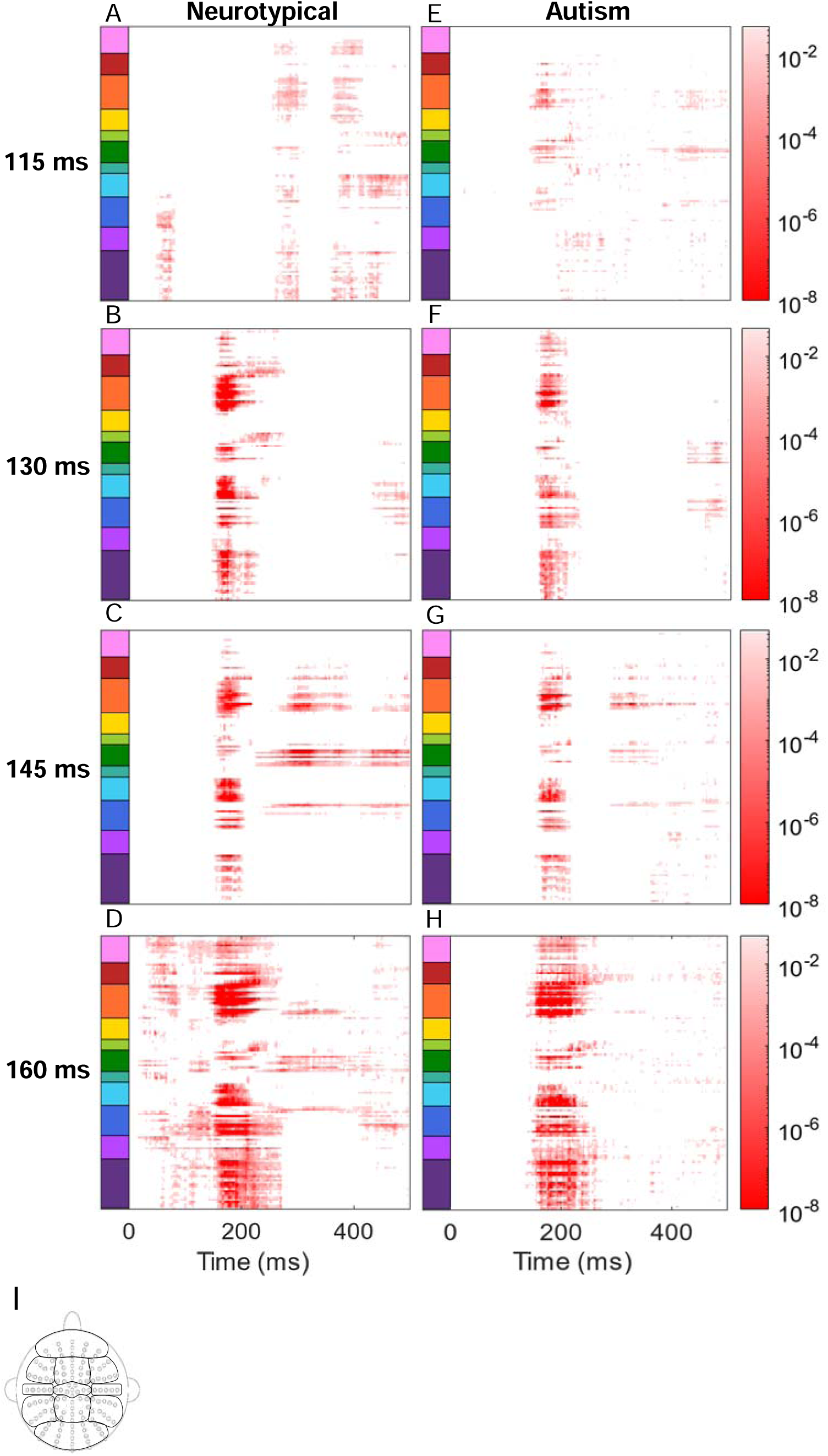
Statistical cluster plots representing the time and location of significant ERP amplitude differences between standard and deviant conditions. P-values < 0.05 are in red p-values > 0.05 are in white for the neurotypical (**A.** 115, **B.** 130, **C.** 145, and **D.** 160 ms) and autism (**E.** 115, F. 130, G. 145, and H. 160 ms) groups. I) The y-axis represents clusters of electrodes sorted by location displayed to the left.

### N1

With regard to ‘All Standard’ N1 amplitude, the autism group response was smaller in magnitude than the neurotypical group response (t(58) = -1.99, *p* = 0.05, d = 0.49). Visualization of these differences can be seen in the cluster plot in **Figure 6A**, and group average waveforms can be found in **Figure 6B**. Means can be found in **Table 4**.

**Figure 6.**
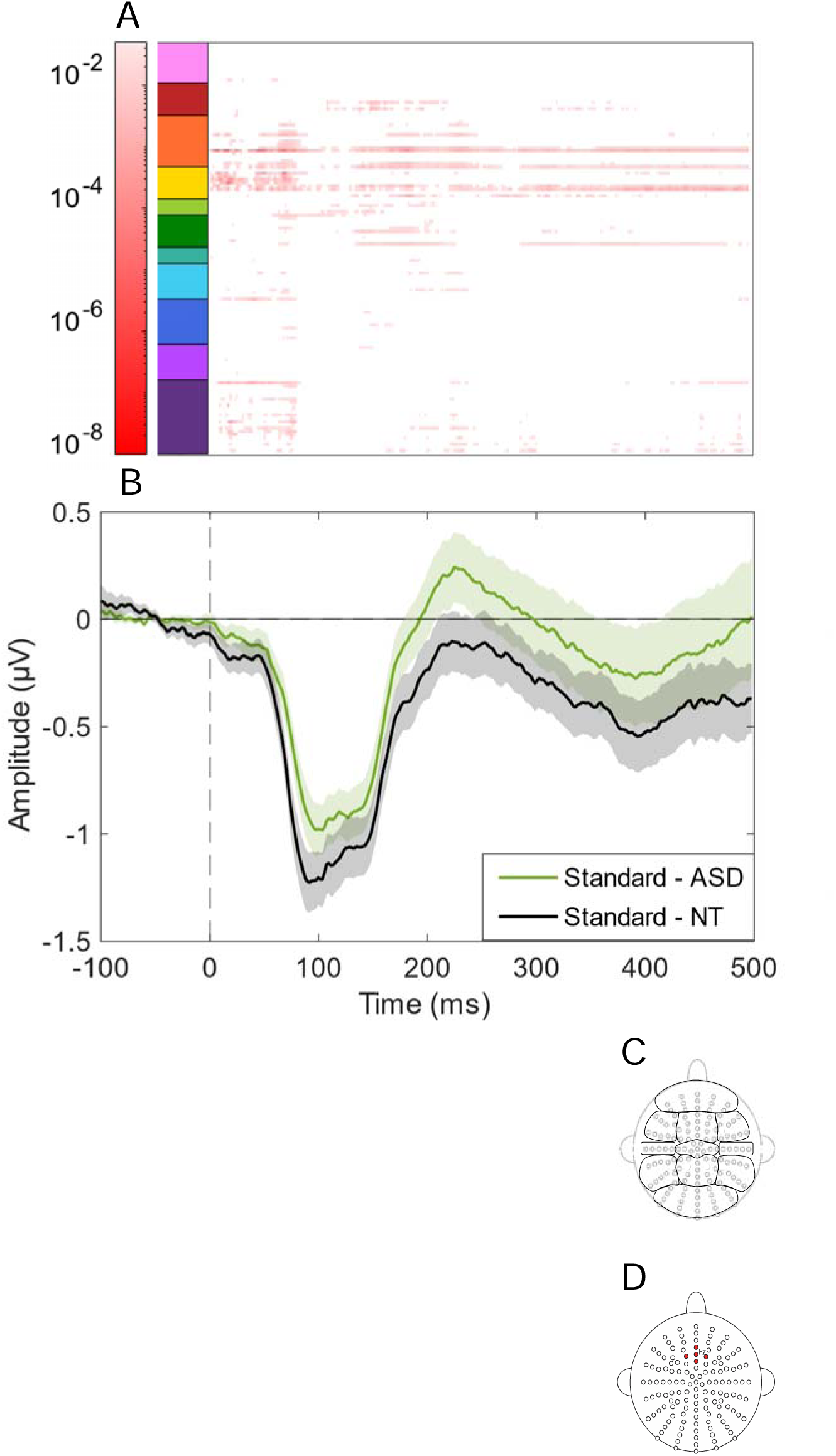
Group-averaged statistical cluster plots and somatosensory evoked potentials (SEPs) comparing the standard vibrations between groups. **A)** Statistical cluster plot comparing ‘All Standard’ p-values (p < 0.05 in red, p > 0.05 in white) at all time points and electrodes between neurotypical and autism groups. **B)** SEP responses to the 100 ms standard vibrations are shown for neurotypical (NT: black trace) versus ASD (green trace) participants, with standard error of the mean indicated by shading around each waveform. **C)** The y-axis in **A** represents clusters of electrodes sorted by location displayed to the right. **D)** The frontal electrode cluster averaged in ‘All Standard’.

**Table 4.** Mean N1 and P220 amplitudes in response to ‘All Standard’ vibrations.

|  | Amplitude Group Mean (SE) |  |
| --- | --- | --- |
|  | Neurotypical | Autism |
| N1 'All Standard' | -1.15 μV (0.10) | -0.90 μV (0.08) |
| P220 'All Standard' | -0.17 μV (0.08) | 0.19 μV (0.12) |

### P220

With regard to the ‘All Standard’ P220 amplitude, the autism group response was more positive than the neurotypical group response (t(58) = -2.52, *p* = 0.02, d = 0.57). Visualization of these differences can be seen in the cluster plot in **Figure 6A**, and group average waveforms can be found in **Figure 6B**. Means can be found in **Table 4**.

### Correlations

Pearson correlations between experimental (N1 and P220) and clinical measures yielded a significant correlation between the averaged Standard P220 and the SRS-2 total (r = 0.29, *p_uncorrected_* = 0.03, *p_corrected_* = 0.23), but did not survive correction for multiple comparisons and are not discussed further here. All other uncorrected *p*’s > 0.13. See **Supplemental Figure 1C** for the correlation plot.

## DISCUSSION

To the best of our knowledge, the present study was the first to detect a somatosensory MMN response in individuals with autism. Since Kemner’s 1994 study, which failed to detect a somatosensory MMN in autism based on stimulus location on the body, no other published EEG studies have tested a purely sMMN in autism (Kemner et al., 1994). Our results demonstrated not only that a MMN was elicited by the same duration conditions in neurotypical and autistic adults, but that the amplitudes of the MMNs elicited were comparable between groups. These results indicated that adults with autism were able to cortically register duration-deviant stimuli with the same sensitivity as the neurotypical adults, namely that the 130, 145, and 160 ms deviants were distinguishable from the 100 ms standard whereas the 115 ms deviant was not. This accords with the behavioral findings, in which there were no group differences in the ability to discriminate somatosensory deviants from standards based on temporal features.

Replication of this pattern of results not only in a larger sample of neurotypical adults, but also in a cohort of autistic adults, reinforces that the mismatch negativity is a valid measure of low-level somatosensory processing (Isenstein et al., 2023). The consistent differentiation between vibrations that vary by 30, 45, and 60 ms – with the absence of this effect when vibrations differ by only 15 ms – further suggests the limits of temporal discrimination under these parameters lies between 15 and 30 ms. The duplication of these findings in the two cohorts - both behaviorally and electrophysiologically - demonstrates the conserved nature of basic somatosensory temporal processes, even across groups that vary significantly in self-reported measures of tactile symptoms. This consistency suggests that although the visual-tactile temporal binding window may be extended in autism (Greenfield et al., 2015; Ropar et al., 2018), isolated tactile temporal perception is likely unaffected, thus implicating multisensory integration as the irregularity.

This main finding of intact sMMN in autism integrates well into the preexisting auditory MMN literature which despite its apparent inconsistency, largely stratifies by age. Several studies found that the auditory MMN was not altered in autism (Ceponiene et al., 2003; Chien et al., 2018; Knight et al., 2020), particularly in adults (Schwartz et al., 2018). Those that did find reduced MMN amplitude in autism were prominently conducted in younger samples (Chen et al., 2020; Schwartz et al., 2018). This pattern suggests that challenges with differentiating between incongruencies in the sensory memory system may exist in children with autism, but effectively resolve by adulthood.

However, while the differences *between* Standard and Deviant vibrations (i.e. the MMN) were comparable between groups across all four duration conditions, exploratory examination found that the underlying electrophysiological waveforms elicited by the “standard” vibrations differed between groups. Specifically, the autism group demonstrated overall smaller amplitude than the neurotypical group within the N1 time window, but larger amplitude in the following P220 time window. This pattern of altered activity suggests that the neurotypical and autistic electrophysiological profiles in response to vibrotactile stimuli differ in small, but potentially meaningful, ways. Indeed, even subtle changes in the neurocognitive processing of sensory information can have downstream effects on higher-order processes like social functioning that can present across the lifespan (Kawakami & Otsuka, 2021).

Given that additional somatosensory processes like discrimination (Zetler et al., 2019), as well as general multisensory processing (Beker et al., 2018; Crosse et al., 2022; Foxe et al., 2015; Ross et al., 2015; Wakim et al., 2023), have been shown to be most prominently dysregulated in children, but not adults, with autism, it is worth considering that delayed development of sensory-perceptual processes may be responsible. Prior work by our group in neurotypical cohorts showed a trend toward smaller N1 and larger P2 amplitudes in younger ages that receded with age, reminiscent of the current findings in autistic adults (Uppal et al., 2016). Accordingly, atypical SEPs in autism may relate in part to altered developmental trajectories in somatosensory temporal processes. In this case, it might be expected that younger individuals with autism – possibly preceding the divergence of these processes – would have altered sensitivity in temporal processing of somatosensory stimuli and may demonstrate differences in sMMN. Together, these considerations converge on the critical role of cortical maturity in sensory perception, while also highlighting the robustness of certain low-level processes like deviance detection.

### Limitations

While this study maps the sMMN responses in multiple duration deviant conditions, generalized conclusions are limited because we did not counterbalance the duration of the deviant stimuli relative to the standard stimuli by including conditions where the deviant stimuli were shorter than the standard stimuli. Prior work has demonstrated that duration deviants both longer and shorter than the standard can elicit a MMN (Näätänen et al., 1989), including in the somatosensory domain (Spackman et al., 2007), but this avenue warrants further exploration in autism. Further, the present study includes a relatively wide age range spanning young and middle adulthood. As discussed above, the potential implications of age and cortical maturity on the detection of deviant sensory stimuli in autism are substantial and future work should include younger age groups.

Taken together, the outcomes of this study suggest that autistic and neurotypical adults detect duration differences in vibrotactile stimuli in similar ways, both behaviorally and electrophysiologically. However, underlying differences in the ERPs may suggest that somatosensory processing differs fundamentally despite conserved deviance detection.

## Acknowledgments

We would like to acknowledge Ashley Xu for her assembly of the Arduino device and Presentation scripts. We also thank Emma Mantel, Laura Ziemer, Philbert Ndagijimana, Yacinda Hernandez, and Grace Rico for their assistance with data collection. Finally, we thank Leona Oakes for her training and supervision in the administration of the ADOS-2.

## Availability of data and material

The datasets used and/or analyzed during the current study are available from the corresponding author on reasonable request.

## DECLARATIONS

### Ethics approval and consent to participate

All aspects of the research conformed to the tenets outlined in the Declaration of Helsinki, with the exception that this study was not preregistered. The institutional review board of the University of Rochester, where the data collection took place, approved this study (STUDY00002036). All participants, or caregivers as appropriate for adults with assigned legal guardians, provided written informed consent. Adults with assigned legal guardians also provided verbal and written assent to participate.

### Consent for publication

Not applicable

### Author contributions

JJF and EGF conceived the study and designed the original experiment. ELI recruited and phenotyped the participants as well as collected the data. ELI analyzed the data and created the illustrations and wrote the first draft of the paper, in close collaboration with JJF and SM. JJF, SM and EGF provided substantial editorial input and writing on subsequent drafts. All authors read the final draft and provided critical input.

---

**Supplemental Figure 1:**
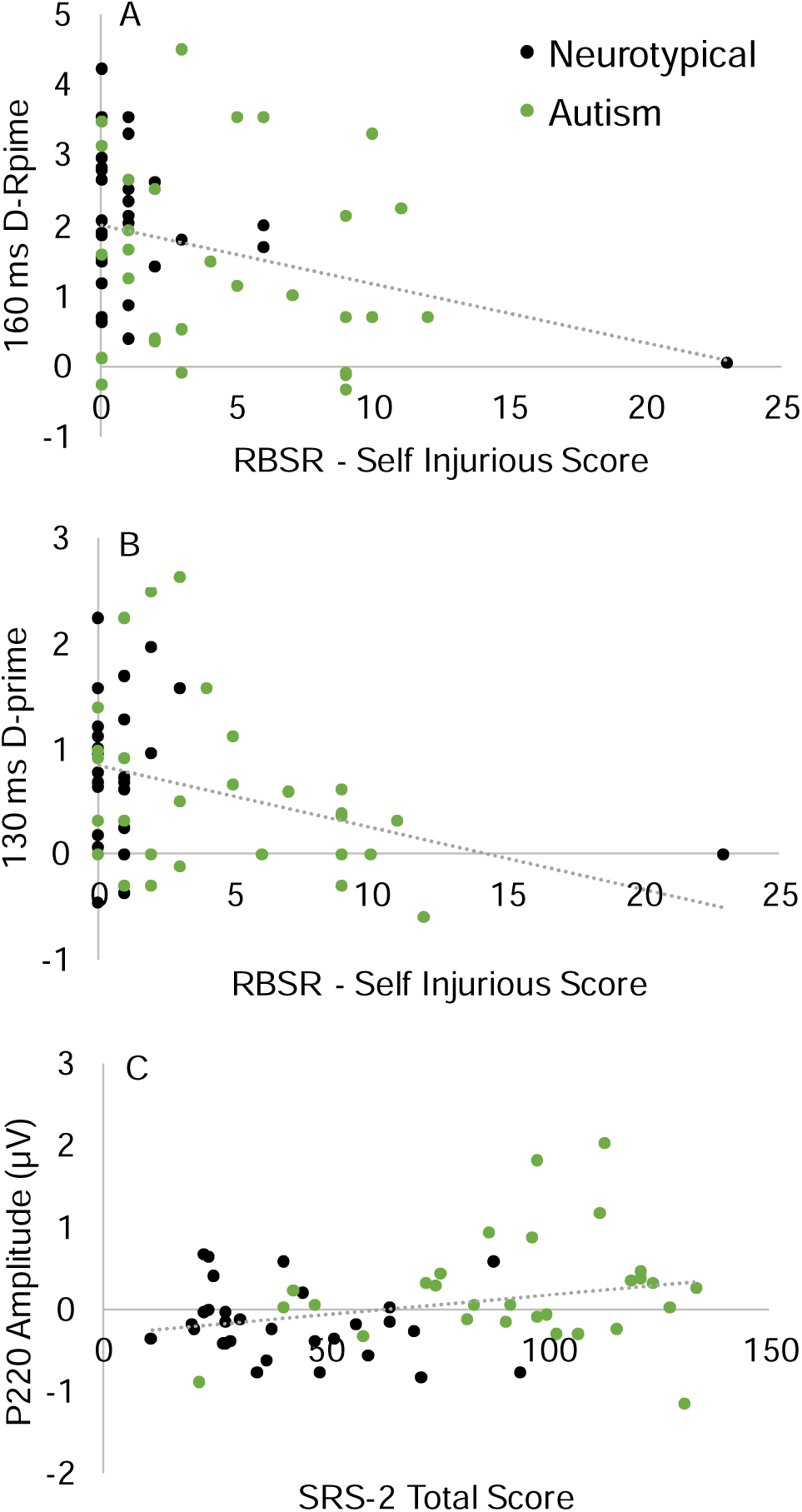
**(A)** Correlation between RBSR - Self Injurious Score and 160 ms D-prime (r = -0.30, p_uncorrected_ = 0.02, p_corrected_ = 0.18). **(B)** Correlation between RBSR - Self Injurious Score and 130 ms D-prime (r = -0.33, p_uncorrected_ = 0.01, p_corrected_ = 0.16). **(C)** Correlation between SRS-2 Total Score and P220 Amplitude (r = 0.29, p_uncorrected_ = 0.03, p_corrected_ = 0.23).

